# Comparing Task-Relevant Information Across Different Methods of Extracting Functional Connectivity

**DOI:** 10.1101/509059

**Authors:** Sophie Benitez Stulz, Andrea Insabato, Gustavo Deco, Matthieu Gilson, Mario Senden

## Abstract

The concept of brain states, functionally relevant large-scale activity patterns, has become popular in neuroimaging. Not all components of such patterns are equally characteristic for each brain state, but machine learning provides a possibility for extracting and comparing the structure of brain states from functional data. However, their characterization in terms of functional connectivity measures varies widely, from cross-correlation to phase coherence, and the idea that different measures provide similar or coherent information is a common assumption made in neuroimaging. Here, we compare the brain state signatures extracted from of phase coherence, pairwise covariance, correlation, regularized covariance and regularized precision for a dataset of subjects performing five different cognitive tasks. In addition, we compare the classification performance in identifying the tasks for each connectivity measure. The measures are evaluated in their ability to discriminate the five tasks with two types of cross-validation: within-subject cross-validation, which reflects the stability of the signature over time; and between-subject cross-validation, which aims at extracting signatures that generalize across subjects. Secondly, we compare the informative features (connections or links between brain regions/areas) across measures to test the assumption that similar information is obtained about brain state signatures from different connectivity measures. In our results, the different types of cross-validation give different classification performance and emphasize that functional connectivity measures on fMRI require observation windows of sufficient duration. Furthermore, we find that informative links for the classification, meaning changes between tasks that are consistent across subjects, are entirely uncorrelated between BOLD correlations and covariances. These results indicate that the corresponding FC signature can strongly differ across FC methods used and that interpretation is subject to caution in terms of subnetworks related to a task.

## 1. Introduction

At a macroscopic level the brain may be conceived of as a collection of specialized regions engaging in dynamic, interactive behavior, typically characterized by the formation of transient functional networks (Bullmore & Sporns, 2009). Functional network formation has been hypothesized to reflect information integration required for higher order cognition (Baars, 2005; Deco, Jirsa, & McIntosh, 2011; Ghosh, Rho, Mcintosh, Kötter, & Jirsa, 2008), implying that despite the ongoing variability, there are regularities that may be used as signatures of cognitive states (Cabral, Vidaurre, et al., 2017). This possibility has traditionally been investigated using whole-brain modeling which explains functional interactions observed in functional magnetic resonance imaging (fMRI) data in terms of graph dynamical systems; i.e. in terms of local dynamics and network interactions (Deco, Ponce-Alvarez, et al., 2013; Deco, Jirsa, & McIntosh, 2013; Deco, Jirsa, McIntosh, Sporns, & Kötter, 2009; Deco, Tononi, Boly, & Kringelbach, 2015; Honey et al., 2009; Jirsa, Sporns, Breakspear, Deco, & McIntosh, 2010). For instance, one such study has recently shown that brain regions exhibit different local dynamics depending on whether human subjects were resting or engaged in cognitively demanding tasks (Senden, Reuter, van den Heuvel, Goebel, & Deco, 2017). Other whole-brain modeling studies have shown that network interactions likewise exhibit differences across cognitive states (Pallares et al., 2018; Senden et al., 2018). Lately, machine learning has gained popularity as a valuable approach to study cognitively relevant signatures in functional connectivity and assess their generalization capabilities (Abraham et al., 2017; Pallares et al., 2018; Rahim, Thirion, Bzdok, Buvat, & Varoquaux, 2017; Varoquaux et al., 2017; Xie et al., 2017). For instance, recent studies have used machine learning to classify five different cognitive states based on dynamic functional connectivity (dFC) computed over short time windows (Gonzalez-Castillo et al., 2015; Xie et al., 2017). In another study, researchers showed that subjects with schizophrenia exhibit weaker resting-state FC, but with more diversity in functional connections as compared to neurotypical subjects (Lynall et al., 2010). Classification approaches such as these enable the extraction of the most relevant features which, with respect to functional connectivity, constitute those pairwise interactions that are specific to a given cognitive state; that is, its FC signature in the context of other cognitive states in the dataset.

The neuroimaging literature comprises many studies that utilize the multivariate nature of functional connectivity to investigate cognitive processing (Cabral, Vidaurre, et al., 2017; Gonzalez-Castillo et al., 2015; Senden et al., 2018, 2017). This multitude of studies is accompanied by a multitude of choices with respect to how functional connectivity between pre-defined regions of interest (ROIs) may be quantified. Pairwise interactions between ROIs are frequently measured in terms of the Pearson correlation (Deco & Jirsa, 2012; Deco, Senden, & Jirsa, 2012; Du, Fu, & Calhoun, 2018; Honey et al., 2009; Jangraw et al., 2018) or covariance (Cole, Yang, Murray, Repovš, & Anticevic, 2016; Gilson, Moreno-Bote, Ponce-Alvarez, Ritter, & Deco, 2016) between their respective signals. However, some studies have instead used pairwise partial correlations in the form of the inverse of the regularized covariance matrix (Smith et al., 2011; Varoquaux, Gramfort, Poline, & Thirion, 2010). We also note that inverse covariance is the connectivity parameter of a Gaussian graphical model and can thereby be considered an undirected effective connectivity measure (Gilson et al., 2019). Nevertheless, we will refer to all connectivity measures here as FC. When it comes to dynamic functional connectivity, the range of choices is even broader. One might, for instance, choose to compute dFC using a sliding-window approach (Cabral, Kringelbach, & Deco, 2017; Hutchison et al., 2013). In this case options regarding FC metric (Pearson correlation, covariance etc.) need to be considered in light of choices regarding window length and vice versa. Additionally, dFC offers further choices for metrics in the form of instantaneous phase coherence (iPC; Cabral, Kringelbach, et al., 2017; Glerean, Salmi, Lahnakoski, Jääskeläinen, & Sams, 2012; Senden et al., 2014) and the outer product of the leading Eigenvector of the iPC matrix (Cabral, Vidaurre, et al., 2017). Note that the latter two do not require sliding windows.

Considering this wealth of choices, the question arises to what extent they produce consistent results with respect to FC signatures of cognitive states. The goal of the present study is to address this open issue. This requires in the first instance to evaluate to what extent different FC metrics are suitable for classification and, more importantly, how well classification results generalize in a rigorous cross-validation procedure. For if an FC metric fails to exhibit generalizable classification performance, it is unlikely to produce consistent FC signatures. Therefore, we employ machine learning to evaluate to what extent choice with respect to metric (for both static and dynamic FC) and time-scale (for dynamic FC) affect classification among five cognitive states as well as the identification of FC signatures of these states. For machine learning techniques, the choice among several types of cross-validation can reveal different aspects of the regularity/variability in the data. Here, evaluation was conducted using two types of cross-validation: Within-subject cross-validation, which focuses on time scales in the experiment (e.g. with succession of blocks) quantifying at information that generalizes over time; and between-subject cross-validation, which aims at extracting information that generalizes across subjects. The states include rest and performance of one of four tasks: an n-back task, the Eriksen flanker task, a mental rotation task, and an odd-man-out task (cf. Senden et al., 2018, 2017). We find that both the choice of FC metric and time-scale strongly impact FC signatures. However, the choice of FC metric is especially consequential since different metrics may produce completely distinct signatures. This calls for a more careful approach towards the interpretation of results stemming from different definitions of functional connectivity.

## 2. Material and methods

### 2.1 Functional Data

We used resting and task-state blood-oxygen-level-dependent (BOLD) time-series data from fourteen healthy subjects (eight females, mean age = 28.76) previously described by Senden et al. (2017). Briefly, data originate from a single MR scan session comprising five functional runs. These runs included a resting-state measurement (eyes closed) and four individual task measurements. The latter consisted of a visual n-back task (Kirchner, 1958), the Eriksen flanker task (Eriksen & Eriksen, 1974), a mental rotation task (Shepard & Metzler, 1971), and a verbal odd-man-out task (Flowers & Robertson, 1985). Each task was continuously performed for a period of 384 seconds (192 2-second volumes). The resting state run comprised the same number of data points as the tasks. For the n-back and flanker task stimuli were presented at a rate of 0.5 Hz; for the mental rotation and odd-man out tasks they were presented at a rate of 0.25 Hz. Task sequence was counterbalanced across participants with the exception that the resting state functional run was always acquired first to prevent carry-over effects (Grigg & Grady, 2010). The data were acquired using a 3 Tesla Siemens Prisma Fit (upgraded Tim Trio) scanner and a 64-channel head coil. Anatomical images were automatically processed with the longitudinal stream in FreeSurfer (Reuter, Schmansky, Rosas, & Fischl, 2012; http://surfer.nmr.mgh.harvard.edu/) including probabilistic atlas based cortical parcellation according to the Desikan_Killany (DK) atlas (Desikan et al., 2006). Initial preprocessing of functional images including slice scan time correction, 3D-motion correction, high-pass filtering with a frequency cutoff of 0.01 Hz and registration of functional and anatomical images was performed using BrainVoyager QX (v2.6; Brain Innovation, Maastricht, the Netherlands). The remaining processing steps were performed in MATLAB (version 8.10.0. Natick, Massachusetts: The MathWorks Inc., 2013). Specifically, signals were cleaned further by performing wavelet de-spiking (Patel & Bullmore, 2016) and regressing out a global noise signal given by the first principal component of signals observed in the ventricles. Following this, the average BOLD signal for each of 68 cortical regions defined by the DK atlas was computed.

### 2.2 Spatiotemporal functional connectivity

#### 2.2.1 Blood-Oxygen-Level-Dependent Signal

While the BOLD signal does not belong to the category of FC measures we include it in the analysis to underline the merit of using second-order metrics such as FC. Here we use the pre-processed timeseries of the BOLD signal of each region of interest (ROI).

#### 2.2.2 Covariance

Dynamic covariance (dCov) between regions *i* and *j* within a time window *w* was calculated as follows:

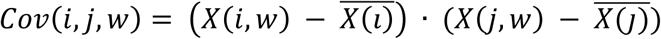

where, *X*(*k*, *w* ^)^ is the BOLD signal in region *k* observed for time window *w* and 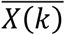 is the mean BOLD activation in region *k*. We calculated dCov for window lengths of 20, 40, 60, 80, 100 and 120 seconds with a timestep of 4 seconds. We also computed the static covariance matrix (sCov) by extending the window to cover the full signal.

#### 2.2.3 Pearson Correlation

Dynamic Pearson correlation (dPC) between regions *i* and *j* within a time window *w* was calculated as follows:

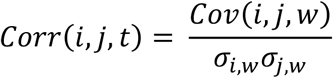

where, σ_*k*, *w*_ is the variance of the BOLD signal of region *k* within the time window *w*. We calculated dPC for window lengths of 20, 40, 60, 80, 100 and 120 seconds with a timestep of 4 seconds. As for the covariance metric, we computed the static PC matrix by extending the window over the whole session.

#### 2.2.4 Regularized Precision and Covariance

We obtained partial correlation between ROIs as well as regularized precision (inverse of covariance) matrix. To that end, we employed the graphical lasso algorithm (Friedman, Hastie, & Tibshirani, 2008). This algorithm enforces sparsity on the estimation of the precision matrix by imposing an L1 penalty. More specifically, the graphical lasso algorithm minimizes the following function to estimate the precision matrix **K** from the corresponding empirical covariance matrix **S**.

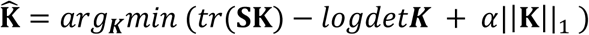

where *α* is the L1 penalty parameter, which is automatically estimated using cross-validation. The estimated covariance matrix is the inverse of the precision matrix. We calculated dynamic precision (dPrec) for window lengths of 20, 40, 60, 80, 100 and 120 seconds with a timestep of 4 seconds.

#### 2.2.5 Phase Coherence

We performed a Hilbert transformation for the BOLD signal of each ROI to obtain its analytical signal. Subsequently, for each ROI pair (i,j), we computed their instantaneous phase coherence (iPC) at a time *t* by taking the cosine of their phase difference:

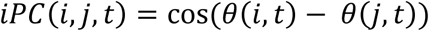

Since this measure is susceptible to high frequency noise, Cabral, Vidaurre, et al. (2017) suggested to compute the outer product of the leading Eigenvector of the iPC matrix at each time step as a more robust measure of instantaneous phase coherence:

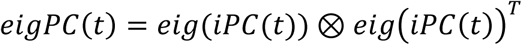

where eig(iPC(t)) denotes the leading eigenvector of the iPC matrix at time *t*.

### 2.3 Classification

#### 2.3.1 Multinomial logistic regression

We use multinomial logistic regression (MLR) as classifier and cross-entropy as our measure of loss. We were interested in how different choices with respect to regularization affect classification performance for different FC metrics. To that end we performed classification either with L1 or L2 regularization. For L1 regularization we use the SAGA algorithm solver (Defazio, Bach, & Lacoste-Julien, 2014), whereas we used a limited-memory Broyden-Fletcher-Goldfarb-Shannon algorithm solver (Bishop, 2006) for L2 regularization. In both cases, we optimized the penalty parameter with nested cross-validation meaning that the parameters were first optimized using cross-validation within the training set before being applied to the entire training set. The classification problem the MLR was trained on included five different classes: The subject at rest, during a visual n-back (n = 2) task (Kirchner, 1958), during the Eriksen flanker task (Eriksen & Eriksen, 1974), a mental rotation task (Shepard & Metzler, 1971), and a verbal odd-man-out task (Flowers & Robertson, 1985).

#### 2.3.2 Cross-validation

##### 2.3.2.1 Within Subject

Due to temporal autocorrelation simple permutation CV produces a test set that is correlated with the training set and thereby does not provide a reliable estimation of classification accuracy for new data. Therefore, we use blocked cross-validation (Roberts et al., 2017). For each task and subject, the samples were divided in 10 consecutive folds. The number of samples contained in each fold depends on the metric chosen. Subsequently, the decoder was trained on the first fold and tested on the second fold. Then the decoder was trained on the first and second fold and tested on the third fold. This procedure was continued until the last fold was reached. The accuracy of the validation procedure was obtained from the mean of the testing accuracy over the 10 trained decoders.

The penalty parameter was optimized using nested cross-validation. More specifically, parameters for each training set were optimized with 2-fold temporal cross-validation on the training set (see figure 2).

##### 2.3.2.2 Between Subject

The decoder was trained on 13 of the 14 subjects and tested on the remaining subject. This procedure was repeated with each subject being left out once. The accuracy of the validation procedure is the median of the testing accuracy over the 14 trained decoders.

The penalty parameter was optimized using nested cross-validation. More specifically, parameters for each training set were optimized with 13-fold subject cross-validation on the training set (see figure 2).

#### 2.3.3 Pipeline

Before classification, the z-score was taken of each FC measure for each sample in each subject (along the feature axis in figure 2). As can be seen in figure 2 the first axis of the FC matrix corresponds to each sample in their temporal order (varied with time-scale of FC measure e.g. window length). The second axis of the FC matrix corresponds to the vectorized dynamic FC in figure 1B meaning the lower triangle of each connectome of all FC measures (without the diagonal elements). The third axis of the FC matrix corresponds to the 14 subjects in the dataset. In the pipeline the FC measures were tested with within-subject cross-validation to quantify how stable learned classes are across time or between-subject cross-validation to quantify how stable learned classes are across subject as described in the previous sections. The classification pipeline was implemented in python using the Scikit-learn library (Pedregosa et al., 2011).

**Figure 1.**
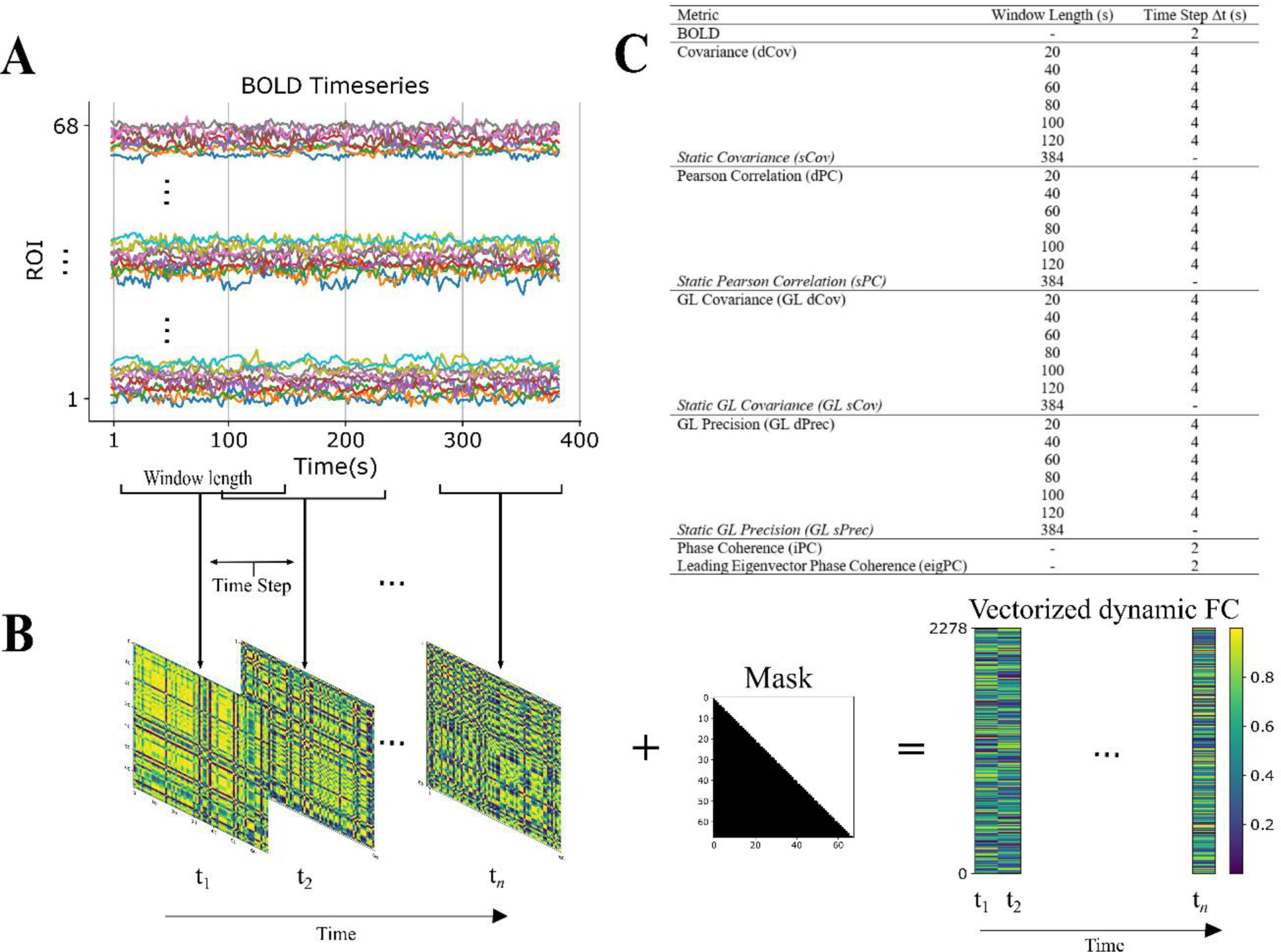
Extracting FC from BOLD signal (A) BOLD signal of 68 ROIs for 384s of an fMRI session. (B) FC matrices extracted from the BOLD signal in windows of length WL. To eliminate duplicate values a mask is applied and (ROI^*^(ROI−1))/2 = 2278 features are obtained for each timepoint t (diagonal elements not included). (C) Table of FC types calculated from the BOLD signal.

**Figure 2:**
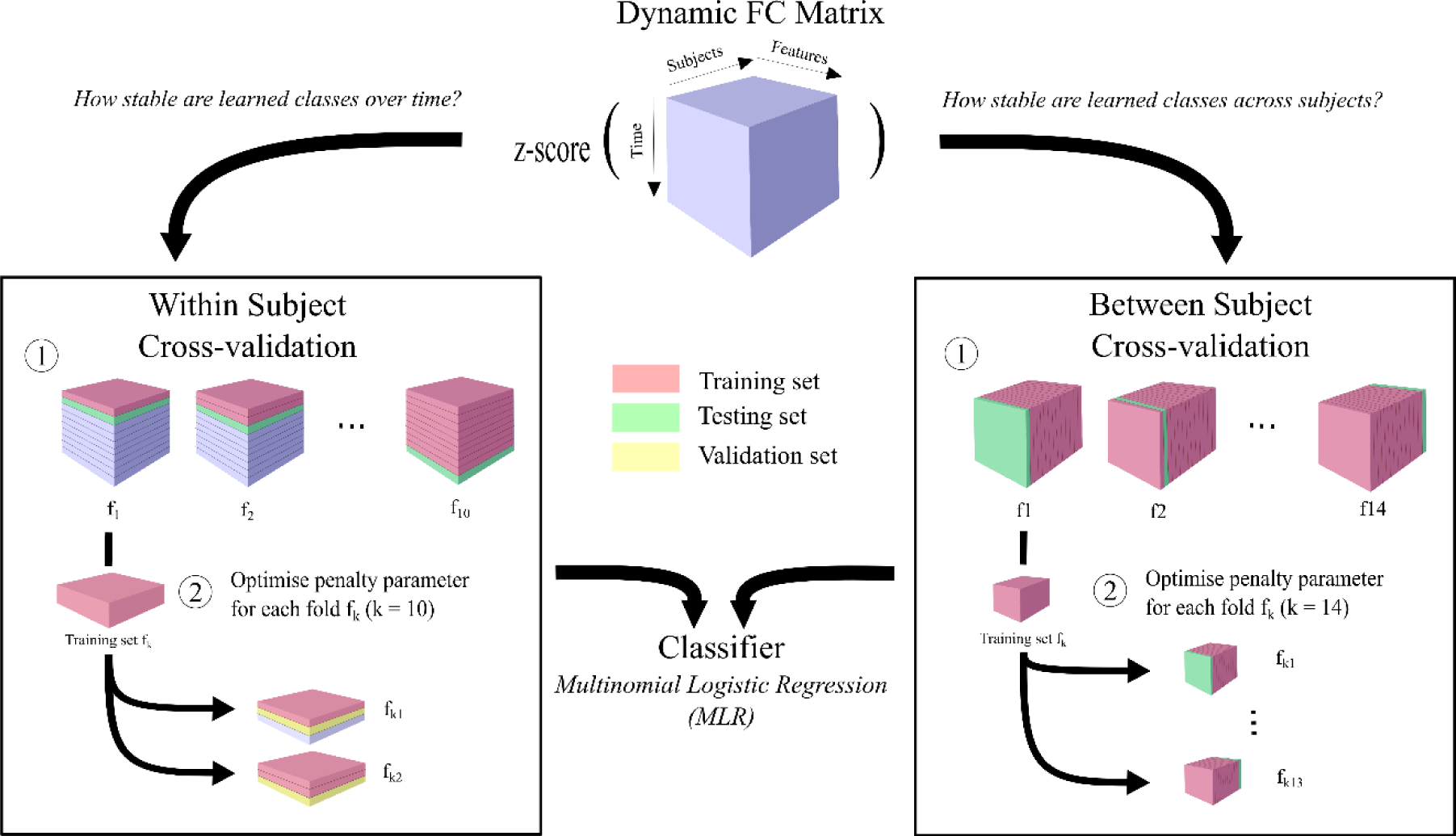
Within- and between-subject cross-validation procedure. In within-subject cross-validation (left) the data is split in sections along time, whereas in between-subject cross-validation (right) the data is split by subject.

#### 2.3.4 Recursive feature elimination

Recursive feature elimination (RFE) iteratively removes the feature that is least important for classification (Guyon, Weston, Barnhill, & Vapnik, 2002). We deemed those features leading to maximum accuracy using temporal or subject cross-validation as the best features to use for classification. The ranking of all features obtained by the RFE was indicative of the structure of the information obtained from each FC; i.e. the state signature. The number of best features was also selected within the nested cross-validation before optimizing the penalty parameter.

### 2.4 Similarity Measure

We used Spearman Rank correlation *r*_*s*_ to quantify the similarity between selected features (state signatures) when using different FC metrics at different time scales.

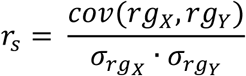

where, *rg*_*X*_, *rg*_*Y*_ respectively are the ranks of variables X and Y, *cov*(*rg*_*X*_, *rg*_*Y*_) is the covariance of the rank variables and σ_*rgX*_, σ_*rgY*_are their standard deviations.

## 3. Results

### 3.1 Classification Performance of FC metrics

#### 3.1.1 Covariance

We first examined the classification performance of dynamic functional connectivity measured in terms of covariance for the classification of rest and four different tasks. For within-subject classifications, median accuracy increases monotonically with window length starting at 68 % correct for a 20 second window and plateauing at perfect classification for windows longer than 80 seconds (figure 3A).

**Figure 3:**
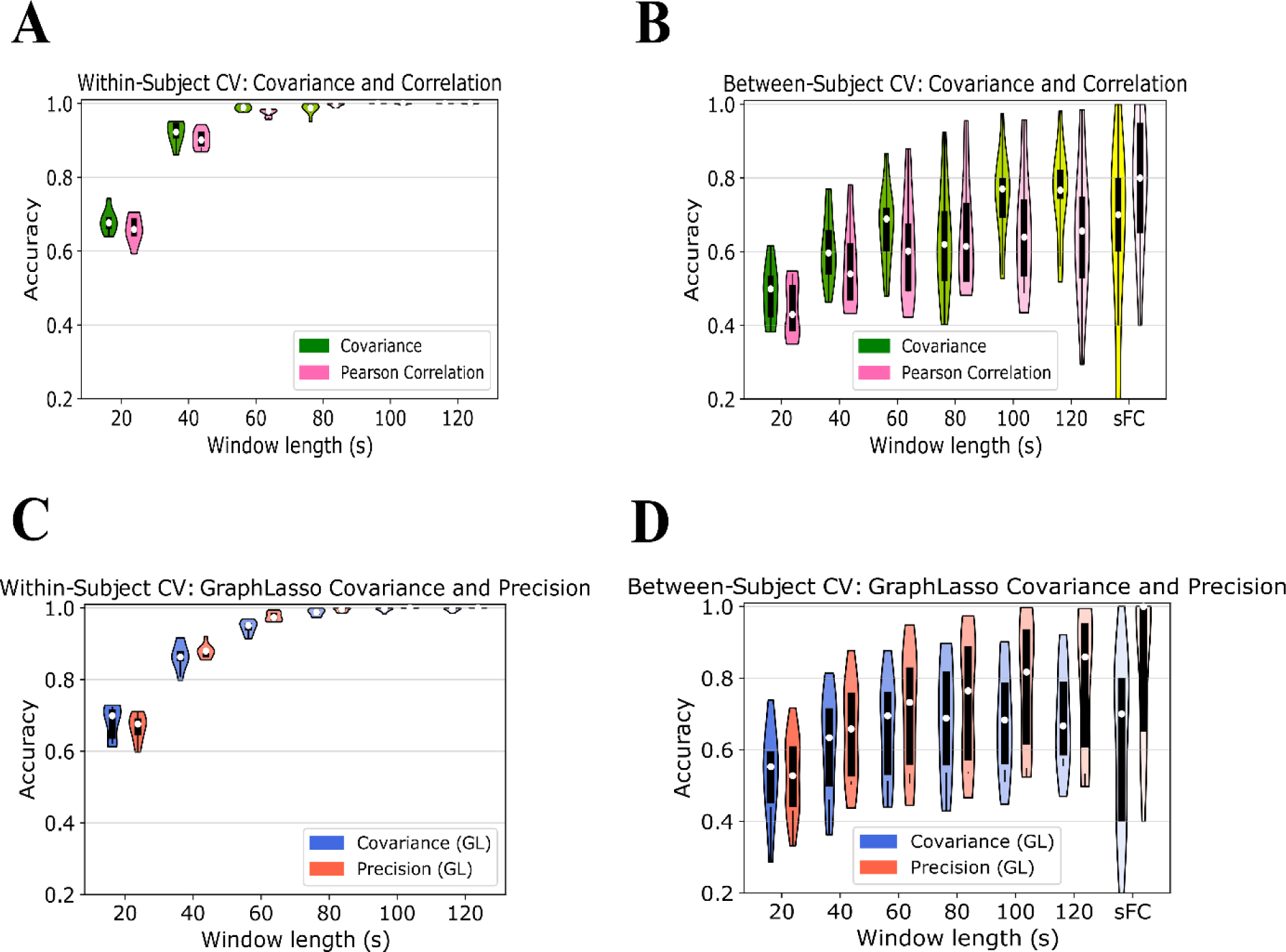
Performance of regularized FC measures GraphLasso covariance and precision. (A) Within-subject cross-validation accuracy using GraphLasso covariance (green) and GraphLasso precision (pink). (B) Between-subject cross-validation accuracy using GraphLasso covariance (green) and GraphLasso precision (pink). Chance level is 0.2.

For the between-subject classification task, median accuracy increases non-monotonically with window length. Specifically, while accuracy generally increases with window length, plateauing at 77% correct for a window length of 100 seconds, it exhibits a dip at 80 s (figure 3B). It is possible that the frequency bands captured by a window length of 80 s reflect processes common to task execution while those captured by a window length of 60 s do not. As such the dFC patterns at 80 s would become more similar which may decrease the signal-to-noise ratio. More similar patterns are difficult to distinguish and can lead to a decrease in classification performance. Static covariance achieves an accuracy of 70 %, similar to what can be achieved with a window length of 60 seconds. At large window lengths, such as in static covariance, covariance may be dominated by lower frequency bands which do not differ strongly across tasks. These could mask more task-relevant, high frequency bands as captured with window lengths of 120 s.

These results show that states signatures are easily extracted in within-subject CV, but are severely affected by subject and run-specific variability in between-subject CV.

#### 3.1.2 Pearson correlation

Next, we examined the classification performance of dynamic functional connectivity measured in terms of Pearson correlation. For the within-subject classification task, median accuracy increases monotonically starting at 66 % correct for a window length of 20 s and saturating at perfect classification for windows longer than 80 s (figure 3A).

For between-subject classifications, median follows a monotonically increasing trend starting at 43 % for a window length of 20 s and reaching 66% for a window length of 120 s (figure 3B). The between-subject median classification accuracy for static Pearson correlation is 80 % which is roughly 15 % higher than median accuracy for a window length of 120 s. This suggests a continuation of the monotonically increasing trend beyond window lengths of 120 s. The trends observed in empirical covariance and correlation suggest that they are differentially affected by noise in the data. At window lengths until 120 s covariance generally performs better, possibly because the standardization in the Pearson correlation also removes information with the variance at shorter time scales. At longer time-scales variance likely contains more noise and its removal increases performance as with static Pearson correlation. Nevertheless, Pearson correlation systematically gives worse results than covariance for each window size.

#### 3.1.3 GraphLasso Covariance

Hereafter, we examined the classification performance of dynamic functional connectivity measured with covariance obtained from the graphical lasso estimation. Within-subject median classification accuracy of GraphLasso covariance increases monotonically from 93 % at a window length of 20 s and saturates at 100 % with window lengths from 60 s on (figure 3C).

Between-subject median classification accuracy of GraphLasso covariance increases non-monotonically from 55 % at a window length of 20 s, reaches a maximum of 70 % at a window length of 60 s and decreases again to 67 % at a window length of 120 s (figure 3D). The median accuracy increased again to 70 % for static GraphLasso covariance. Similar to empirical covariance, regularized covariance does not improve at longer window lengths, suggesting that it might be affected by noisy fluctuations in the low-frequency range.

#### 3.1.4 GraphLasso Precision

Next, we compared the classification performance of dynamic functional connectivity measured with the regularized precision which underlies the regularized covariance. Within-subject median classification accuracy of the GraphLasso Precision increases monotonically from 88 % at a window length of 20 s and reaching 100 % accuracy at window length beyond 100 s (figure 3B). Regularized as well as empirical metrics follow a comparable trend, suggesting that they are affected by similar noisy temporal fluctuation at shorter time-scales.

Between-subject median classification accuracy of GraphLasso precision shows a monotonically growing trend continually increasing from 53 % at a window length of 20 s until 86 % at a window length of 120 s without saturating (figure 3C). The median accuracy of static GraphLasso precision reaches 100% following the monotonically increasing trend. Except for the 20s-window, it GraphLasso precision gives the best performance compared to all three measures considered. This suggests, that removing noisy fluctuation by estimating the underlying precision is preferable to standardizing by the variance as in the Pearson correlation.

#### 3.1.5 Phase coherence

Lastly, we examined the classification performance of dynamic functional connectivity measured by phase coherence. The within-subject median classification accuracy was 19 % for the BOLD signal, 42 % for phase coherence, 32 % for the leading Eigenvector of phase coherence and 66 % for the Pearson correlation with a window length of 20 s (figure 4A).

**Figure 4:**
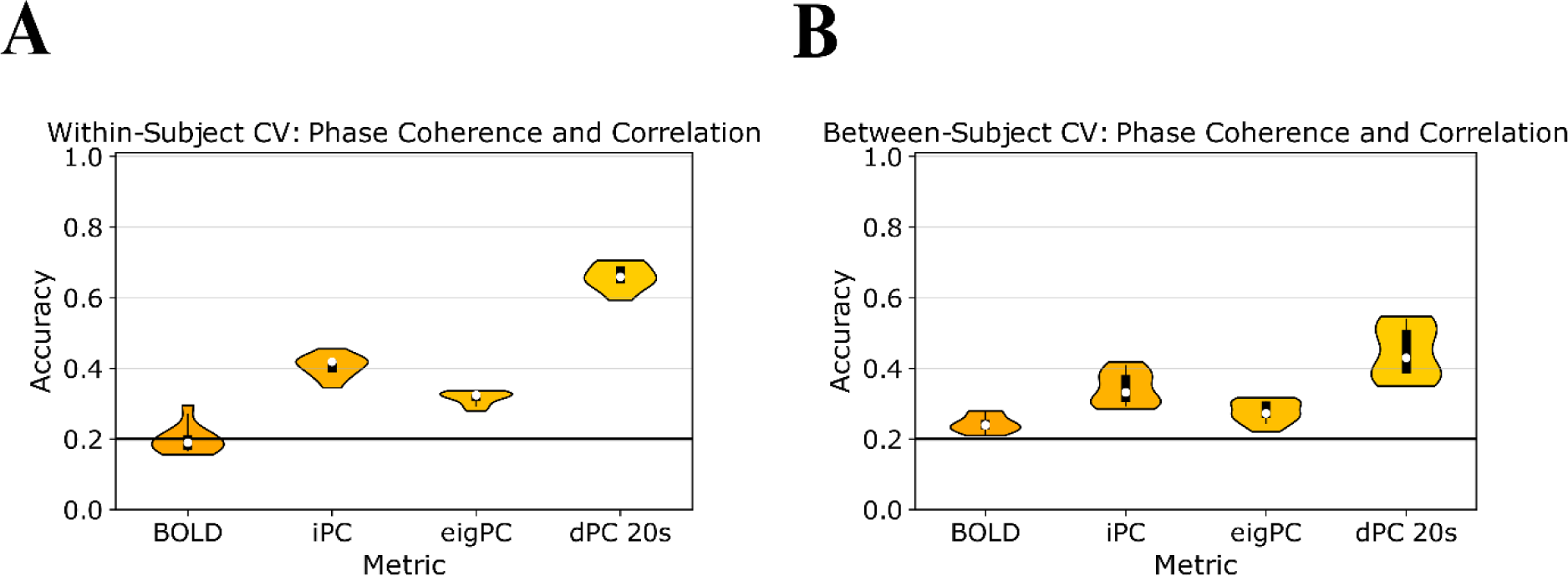
CV Accuracy at short time-scales. (A) Within-subject CV accuracy of the BOLD timeseries, phase coherence (iPC), the leading Eigenvector of the phase coherence (iPC) and Pearson Correlation with a window length or 20s and a time step of 4 s (dPC 20s). (B) Between-subject CV accuracy of the BOLD timeseries, phase coherence (iPC), the leading Eigenvector of the phase coherence (iPC) and Pearson Correlation with a window length or 20s and a time step of 4 s (dPC 20s). Chance level (0.2) indicated with black line.

The between-subject median classification accuracy was 24 % for the BOLD signal, 33 % for phase coherence, 27 % for the leading Eigenvector of phase coherence and 43 % for the Pearson correlation with a window length of 20 s (figure 4B). Although phase coherence has been found to resemble Pearson correlation with a window length of 20 s (Pedersen, Omidvarnia, Zalesky, & Jackson, 2018), it appears to retain less task-dependent information content than the Pearson correlation. Pearson correlation performs ~ 20 % better in within-subject classification and ~10 % better in between-subject accuracy. Interestingly, the leading Eigenvector of phase coherence (eigPC) performs worse than phase coherence suggesting that more variability is desirable and that this method not only remove noise, but also temporally stable information. However, one should keep in mind that the goal of eigPC was to extract the expression of microstates within brain states, whereas this paper aims to quantify the stationarity of state signatures. Our results suggest, that microstates may individually carry only limited state specific information, but their distribution might be more state specific as brain states can be distinguished beyond chance level. Considering the 20 % chance level, phase coherence performs poorly for both within- and between-subject classification, although it outperforms the BOLD signal, where the task-dependent information does not seem as accessible as with dFC. This underlines the need of second-order metrics such as functional connectivity in studying brain functions.

### 3.2 Effects of Regularization

Lastly, we investigated the effect of regularization on classification performance. Regularization is a commonly used tool to prevent a classifier from overfitting the training data (Bishop, 2006). Using L1 regularization instead of L2 regularization in our classification did not improve the performance of the classifier. Rather it reduced accuracies by approximately 3% on average (see S2), suggesting that the information is distributed across functional links, which can be better captured by the L2 regularization.

Another tool that can be used to further reduce dimensionality is feature selection. Here we show the effect of reducing the feature dimensionality of dynamic functional connectivity measured in terms of covariance. For within-subject classification, median accuracy increases monotonically with window length starting at 67 % correct for a 20 second window and plateauing at perfect classification for windows longer than 100 seconds (figure 5B). The median percentage of selected features was 86 % (1957 features) with a window length of 20 s, reached a local maximum at 98 % (2229 features) with a window length of 40 s, and decreased again to 85 % (1827.5 features) with a window length of 60 s. Thereafter, the median percentage increased to 100 % (2278 features) at a window length of 100 s, only to decrease again to 90 % (2060 features) at a window length of 120 s. At no window length did reducing the feature dimensionality significantly increase the classification accuracy.

**Figure 5.**
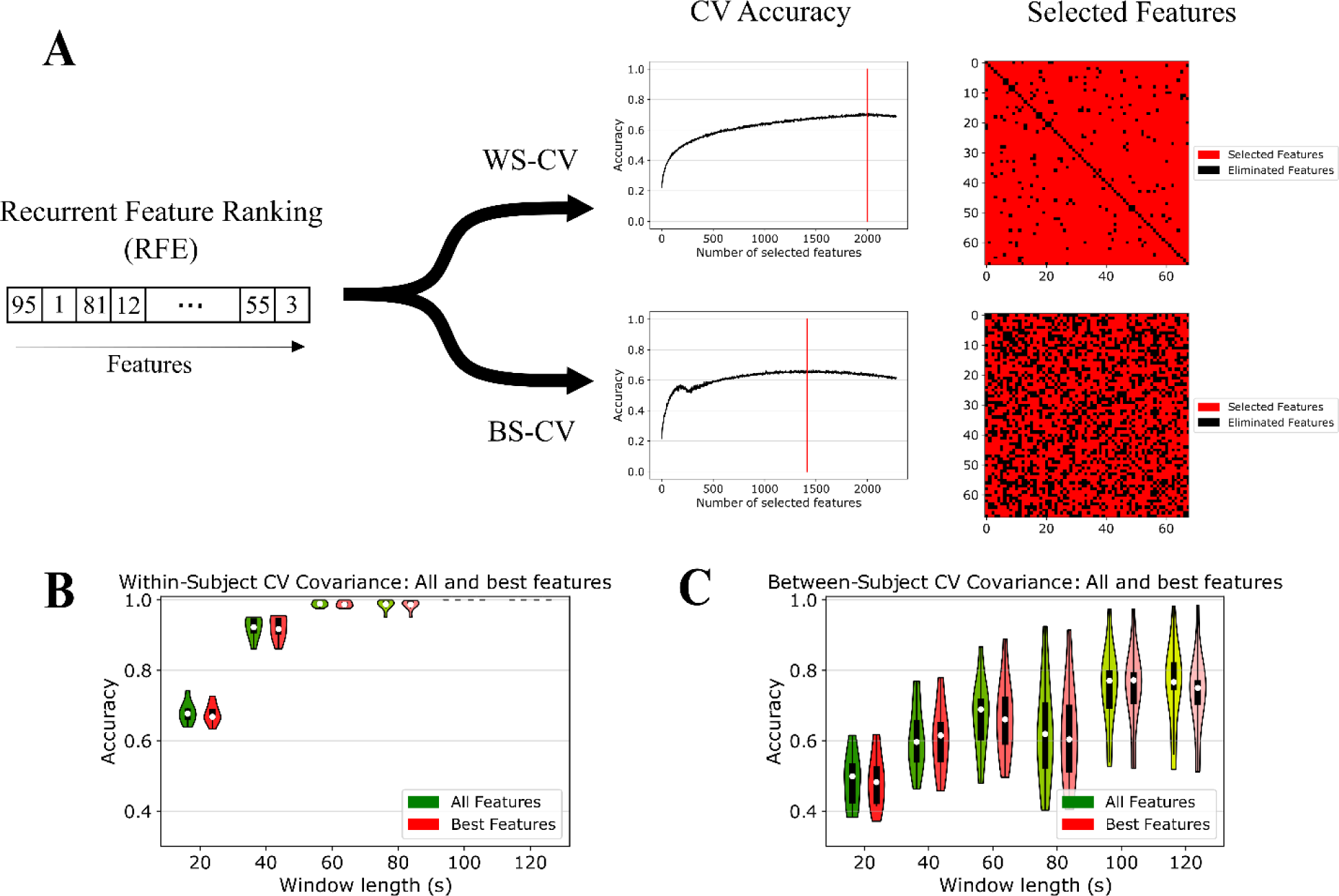
Feature selection performance for covariance. The recurrent feature ranking (RFE) is used to test how many best features give the highest within-subject CV and between-subject CV accuracy. (A) Sketch of RFE procedure. Vertical red line indicates number of features which render the maximal accuracy. (B) Within-subject CV and (C) Between-subject CV accuracy of all features versus best features with covariance. Specific values can be found in the supplementary material (table S5).

For between-subject classification, median accuracy increases non-monotonically with window length starting at 48 % correct for a 20 second window and reaching a maximum at 77 % with a window length of 100 seconds, exhibiting a dip at a window length of 80 seconds at 60 % accuracy (figure 5C). The median percentage of selected features decreased non-monotonically from 84 % (1921.5 features) with a window length of 20 s until 46 % (1060 features) with a window length of 100 s and increases again to 88 % (1994.5 features) with a window length of 120 s. At no window length did reducing the feature dimensionality significantly increase the classification accuracy. The results that neither feature reduction nor using an L1 penalty loss for the classifier lead to an increase in accuracy suggests that the task-dependent information is distributed across features and entangled with the noise in a way that cannot be separated by eliminating features entirely.

### 3.3 Separability among brain states

To identify the origin of the classification errors at different time-scales we plotted the confusion matrices of the within and between-subject classification for covariance. The confusion matrices of within-subject classification only show visible off-diagonal patterns for a window length of 20 s as classification accuracy is high for longer windows. For covariance at a window length of 20 s, rest and the Eriksen flanker task are sometimes confused, the n-back task is sometimes predicted as the Eriksen flanker task, the mental rotation task, or the odd-man-out task, and the mental rotation task is sometimes predicted as the n-back task or the Eriksen flanker task. The confusion matrices of between-subject classification show that rest and the Eriksen flanker task are often confused and that the n-back task and the mental rotation task are also confused. Interestingly, a number of subjects reported that they perceived the n-back task and the mental rotation task as rather effortful, whereas they perceived the Eriksen flanker task as comparatively easy. This suggests that FC may carry signatures of cognitive effort or attention rather than task-specific cognitive processes. The brain during performing a demanding task and the brain at rest could thus correspond to extremes on a spectrum. However, a systematic investigation to what extent task difficulty can be identified based on FC would be required before lending credence to this hypothesis.

The decreasing separability between state-specific signatures at shorter time-scales is likely due to a larger influence of noise on FC estimation. Measures estimated across longer temporal windows are less affected by noise and more stable, as the noise is average out across many timepoints. At shorter window lengths, state-specific signatures are hidden behind noise which is averaged out to a lesser degree during the estimation of each measure due to fewer timepoints.

### 3.4 The structure of task-relevant information differs strongly across time-scale and method of FC extraction

To evaluate to which extent signatures of cognitive states are consistent across FC methods, we perform recursive feature elimination for each method across the whole dataset and compare the resulting rankings using Spearman rank correlation (figure 7). For each dFC measure among GraphLasso precision, GraphLasso covariance, correlation, and covariance, rankings become more dissimilar as the difference in window length increases. This could suggest that task-relevant information content differs across time-scales. However, the concurrent decrease in accuracy for shorter time-scales is in line with the known fact that sufficiently long window lengths are necessary for stable estimates for covariance or correlation. The decrease in accuracy and dissimilarity is greater between shorter window lengths. This may be due to the increasing effect of noise on the FC estimation, which might lead to feature rankings having a more random and thereby less correlated order. Nevertheless, figure 7 suggests that information is relatively robust across time within measure.

**Figure 6.**
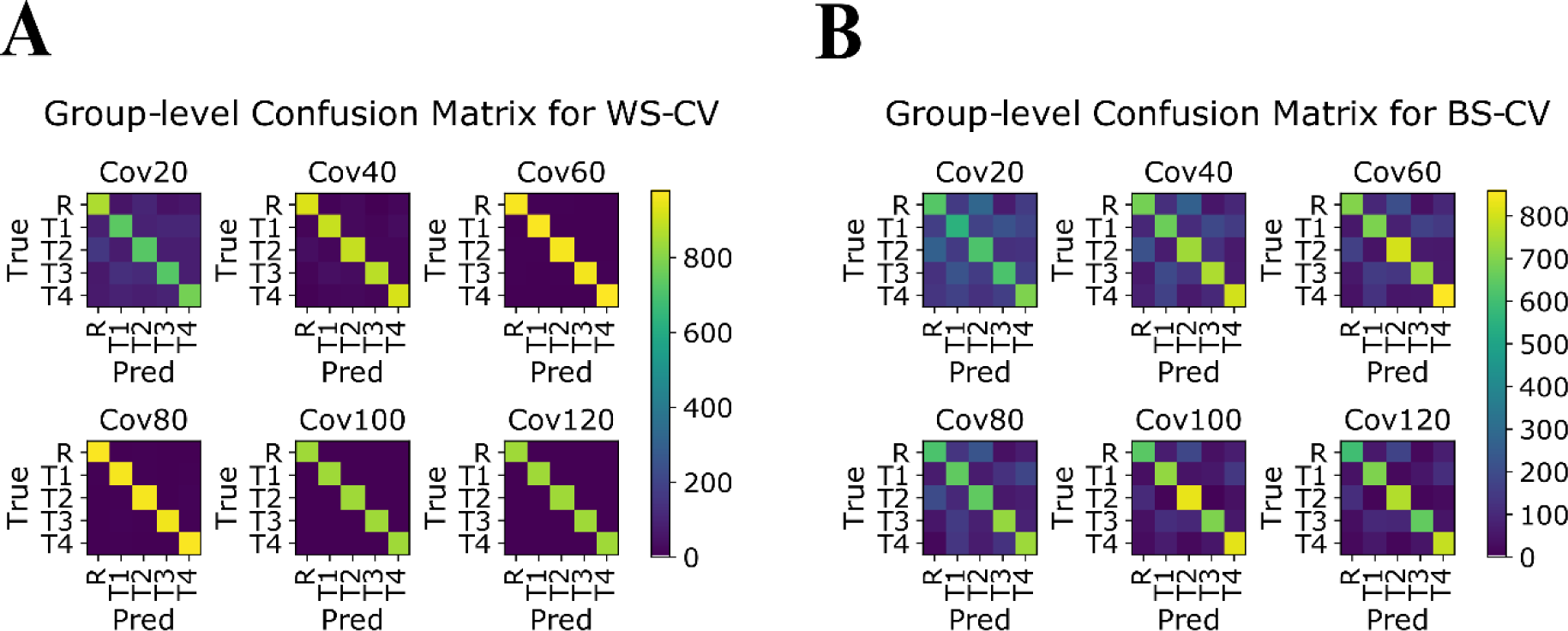
Confusion matrix of covariance for various window lengths. The x-axis indicated the labels predicted by the classifier and the y-axis indicates the true labels. R = rest, T1 = n-back task, T2 = Eriksen flanker task,T3 = mental rotation task, T4 = Odd-man-out task (A) Confusion matrix of the test folds for within-subject cross-validation. (B) Confusion matrix of the test folds for between-subject cross-validation.

**Figure 7.**
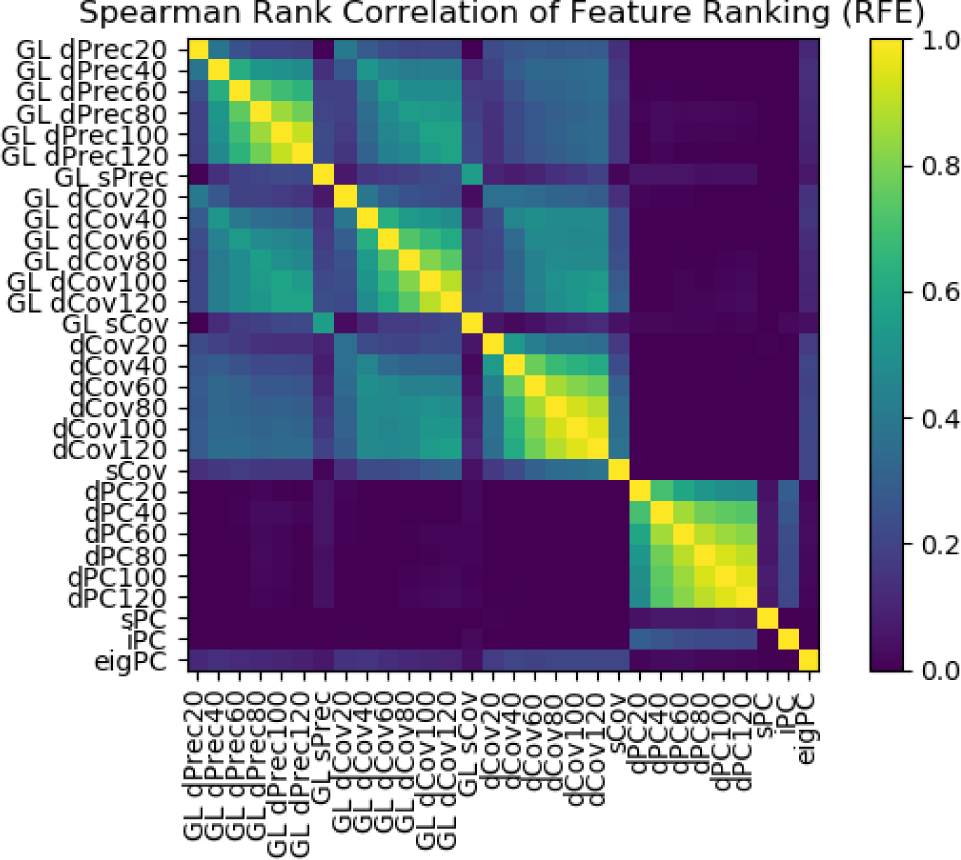
Task-dependent information structure can differ strongly across metric. Spearman rank correlation of all FC metrics.

Surprisingly, covariance and Pearson correlation suggest entirely different features rankings at all time-scales. The dissimilarity is also present between the GraphLasso measures, precision and covariance, and the Pearson correlation. This suggests that the task-relevant information structure (FC signature) retrieved by covariance and Pearson correlation concern distinct pairwise ROI interaction.

At the shortest time-scale, feature rankings obtained from iPC and eigPC do not seem to be correlated even though the measures are very closely related as eigPC corresponds to the axis with the largest variability in iPC. Curiously, the feature ranking in iPC is slightly correlated with the rankings of Pearson correlation metrics and the features ranking of eigPC is slightly correlated with the rankings of covariance metrics.

## 4. Discussion

The aim of this paper was to evaluate the classification performance of FC measures (phase coherence, Pearson correlation, covariance, GraphLasso precision, and GraphLasso covariance) and to what extent the signature of cognitive states extracted from functional connectivity may be sensitive to the choice of FC measures. We use a ground truth in which the four tasks and rest define five distinct brain states.

Firstly, our results show that among non-regularized measures covariance tends to outperform correlation. The ability of regularized measures to accurately capture functional networks can be strongly affected by the length of the window within which it is estimated (Smith et al., 2011). Nevertheless, GraphLasso precision outperformed all measures at all time-scales with the exception of GraphLasso covariance for a window length of 20 s. Regularized metrics capture direct effects between regions and attempt to discard mediating effects from secondary regions on connectivity (e.g. indirect pathways). They are thus more interpretable than measures which do not discard confounding connectivity such as the Pearson correlation or covariance (Varoquaux & Craddock, 2013). Here we have tried two distinct regularization schemes: on the classifier (L2 penalty) and on the measure itself (GraphLasso on each window). We find that the second option is the crucial factor to improve classification accuracy. This supports the choice of regularized approaches (here a graphical model) for extracting connectivity measures that can be used for robust classification.

Secondly, within-subject classification accuracy was generally higher than between-subject classification accuracy and can be conceptualized as an upper limit of accuracy. Within-subject classification accuracy likely reflects temporal stability of cognitive state signatures. Note, however, that in the present dataset, each task was performed in a separate run. This implies that run-specific noise goes hand in hand with state-specific activations and the classifier may to some extent rely on noise to distinguish among cognitive states; i.e. it may distinguish runs. Hence within-subject classification accuracy would likely be lower if each task was performed in multiple distinct runs. Between-subject classification does not suffer from this shortcoming as cognitive states are no longer paired with a specific run. Instead, run-specific variability is accompanied by additional subject-specific variability, rendering classification more demanding. Nevertheless, even the worst performing instantaneous FC measure (eigPC) achieves test accuracies exceeding chance level on this challenging task. A possible reason for the low accuracies in this classification problem may be that the stimuli used here were all visual, making the classification entirely reliant on non-sensory processes. These processes might be hidden by the strong influence of time-, subject- and/or run-specific variability. Therefore, it may be possible that other classification problems reach better accuracies at shorter time-scales and with different FC methods. Classification performance may have been better still if the dataset had included more subjects. As much has already been suggested in a study by Abraham & al. (2017). Furthermore, these authors found that accuracy increases with finer parcellation. The parcellation chosen here was based on pre-defined anatomical landmarks and has been shown to be anatomically valid and reliable across subjects (Desikan et al., 2006).

Thirdly, within- and between-subject classification accuracy increased in proportion with time-scale, which is likely due to high-frequency noise in the signal affecting short time-scales (Cabral, Kringelbach, et al., 2017; Hutchison et al., 2013). An alternative explanation for lower accuracy at shorter time-scales is low task performance (Gonzalez-Castillo et al., 2015). Gonzalez-Castillo et al. (2015) showed that large deviations in task performance are correlated with substantial errors in classification accuracy. These deviations are more likely to bias connectivity measures at shorter time-scales. However, we did not control for this possibility. Interestingly, this would also fit the hypothesis that FC measures carry signatures of cognitive effort rather than task-specific cognitive processes. A second explanation could be that classification depends on specific frequency bands which would require window lengths long enough to capture these functional interactions. This is crucial, since classification assumes a stationary signature exists in each sample. On the other hand, changes in short-timescales may reflect successions of neuronal processes (not just noise) at work within each window. A study investigating the dependence of community structure on window length has already shown that different frequency bands can address distinct neuronal processes (Telesford et al., 2016). Therefore, we need to identify the combination of experimental conditions and extracted measures that exhibit a regularity of patterns, which can be quantified with within-subject classification, that support a good classification. An alternative option is using regularized FC measures aimed at specific frequency bands such as dynamic causal modelling or the Kuramoto model (Cabral, Hugues, Sporns, & Deco, 2011; Friston, Kahan, Biswal, & Razi, 2014). The strong decrease of accuracy at shorter time-scales was driven by the decreasing temporal stationarity as well as a difficulty in distinguishing between specific pairs of tasks and rest, most likely due to noise. This break-down of temporal stationarity was especially visible with a window length of 20 s. Interestingly, the classifier had trouble distinguishing among tasks with similar levels of cognitive effort, which suggests that brain state signatures may carry signatures of cognitive effort or attention rather than task-specific cognitive processes. However, this hypothesis should to be systematically investigated.

Finally, we find that the task-relevant information structure differs far more across connectivity measures than across time-scale. While our results do show that phase coherence is most similar to Pearson correlation with a window length of 20 s as reported in Smith et al. (2011) the similarity between the two is low. This might be due to differences in filtering as the data in the present paper was filtered with a broader band than in Smith et al. (2011). Most importantly, the absence of any similarity in information structure retrieved from correlation and covariance is a counterintuitive result. Correlation is merely normalized covariance and evidence that such closely related methods can provide very different information contradicts the implicit assumption that similar methods should lead to similar conclusions. This is problematic for the interpretation of any results obtained for different measures since there is no ground-truth on task-relevant functional interactions. For example, how would one interpret evidence from studies using network theory to detect communities based on different FC measures (Fuertinger & Simonyan, 2016; Najafi, Mcmenamin, Simon, & Pessoa, 2016; Sporns, 2013)? An encouraging finding is that the task-relevant information retrieved from covariance, GraphLasso precision, and GraphLasso covariance is somewhat similar. This underlines the need for alternative metrics such as regularized FC, where the relationships between the various metrics are better defined (Cabral et al., 2011; Friston et al., 2014; Pallares et al., 2018; Senden et al., 2018, 2017). Within measure the comparability seems to remain, although at a decreasing level as the window lengths become more different. The differences are especially pronounced at lower time-scales and show a similar pattern within each metric. This parallels the decreasing accuracy in within-subject classification which is stronger at shorter time-scales and displays a similar trend for Pearson correlation, covariance, GraphLasso precision, and GraphLasso covariance. Therefore, it is likely that the decreasing similarity along time-scale within measure is due to a breaking down of temporal stability in the signature of brain states. This further underlines the need for sufficiently long window lengths to obtain stable estimates. However, one should also keep in mind that the features used in multinomial classification problems are context-dependent. For example, a feature can be crucial for distinguishing between task A and B, but not between task A and C. The task-relevant information structure extracted by the classifier can thus change depending on the tasks included. Nevertheless, this does not refute our results which falsify the assumption that the same information is obtained from different dynamic functional connectivity measures.

The pipeline developed here can be applied to other neuroimaging tools as well such as electroencephalography (EEG) or functional near-infrared spectroscopy (fNIRS). Quantifying the performance of a classifier is especially important in clinical settings when aiming to identify pathological brain states in new patients. Predictive decoders, for example in the case of brain-computer interfaces, can be implemented with FC metrics, but should be tuned within-subject as the performance is better and more stable.

In conclusion, our findings indicate that the extraction of consistent FC signatures of cognitive processes require sufficiently long window lengths for dynamic functional connectivity estimation and that these signatures are highly sensitive to the choice of functional connectivity measure. This makes it difficult to draw any conclusions on which functional connections are typical for a given cognitive state and calls for greater care in the selection of FC method with respect to the aim of a study as well as more careful interpretation of results stemming from different FC methods.

## Acknowledgements

This research was funded by the European Union’s Horizon 2020 Research and Innovation Programme under Grant Agreements No. 7202070 (HBP SGA1) and 737691 (HBP SGA2). Author GD was also supported by the Spanish Research Project PSI2016-75688-P (AEI/FEDER). The funding source played no role in this study. Furthermore, the authors declare no competing interests.

## Supplementary Material

**Supplementary Figure 1:**
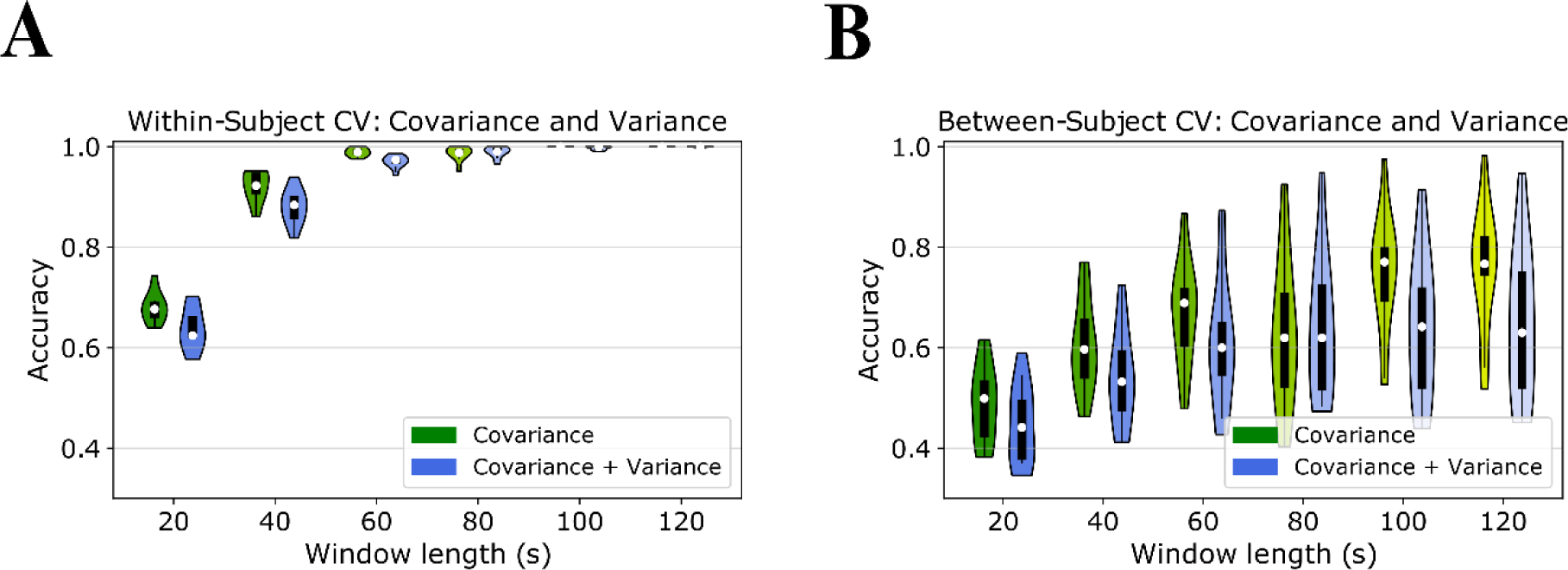
Adding variance to covariance does not outperform covariance alone. (A) Within-subject classification accuracy using only covariance (green) and covariance + variance (blue). (B) Between-subject classification accuracy using only covariance (green) and covariance + variance (blue). Chance level is 0.2.

**Supplementary Figure 2:**
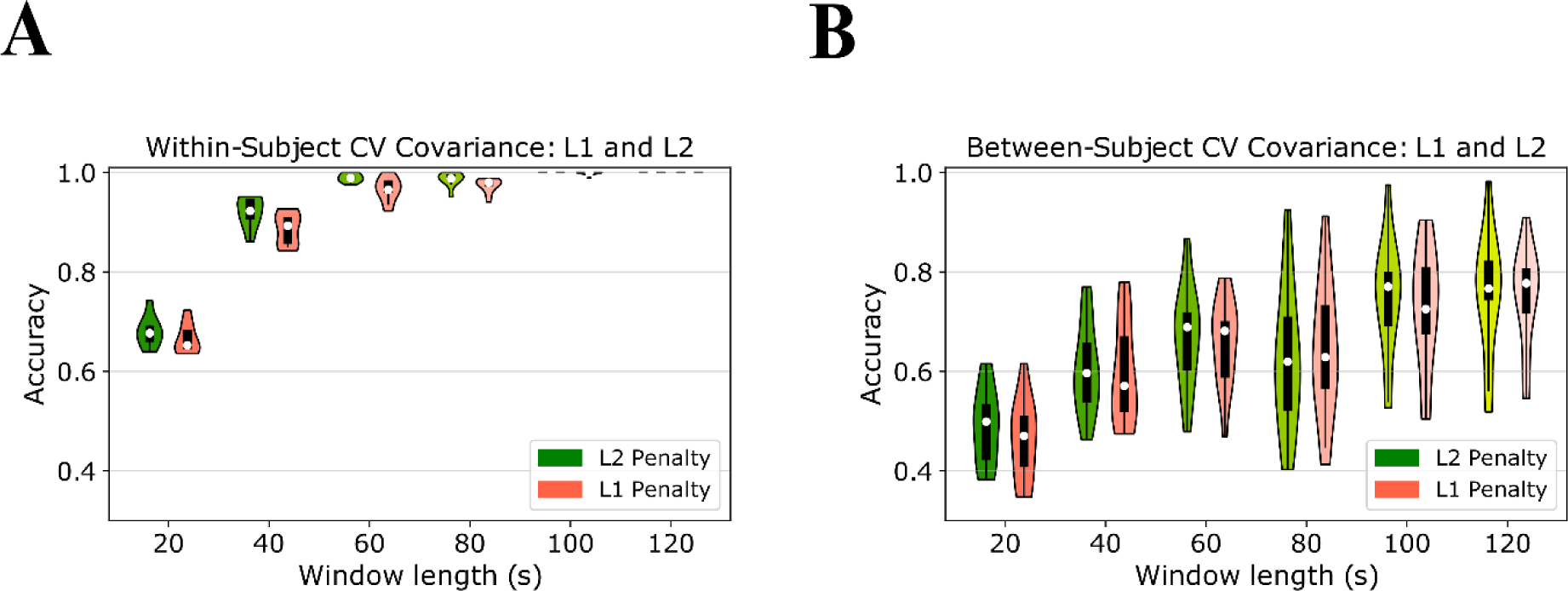
Using L1 regularization instead of L2 regularization does not improve accuracy for covariance. (A) Within-subject accuracy using L2 penalty (green) or L1 penalty (orange). (B) Between-subject classification accuracy using L2 penalty (green) or L1 penalty (orange). Chance level is 0.2.

**Supplementary Figure 3:**
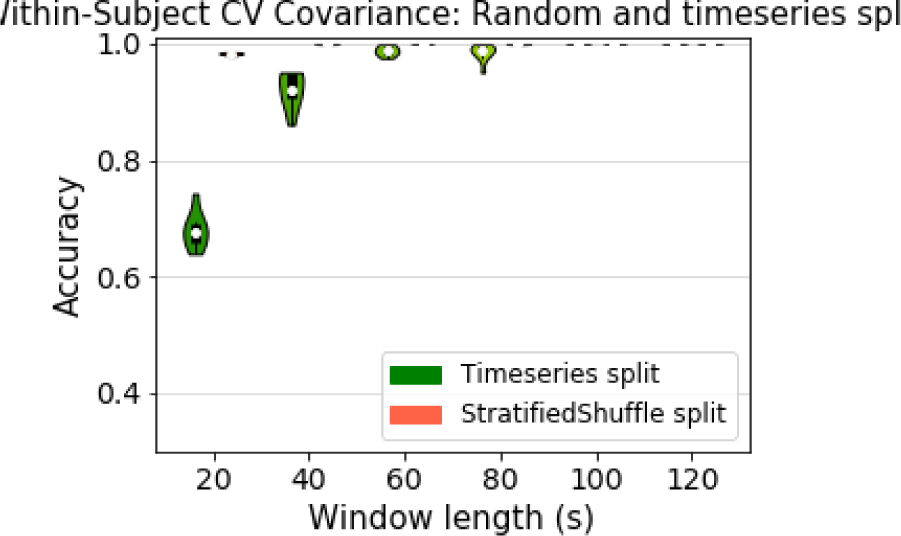
Random split overestimates the cross-validation accuracy within a timeseries. Within-subject classification within a run using covariance with a time series split (green) and a stratified shuffle split (orange). The necessity of within-subject CV to quantify the temporal stability of the classes becomes clear when compared to cross-validation with permutation sets which disregard the temporal autocorrelation (S2). While the permutation CV achieves maximal accuracy for all window lengths, within-subject CV shows a break-down of temporal stability which has also been shown by other studies (Roberts et al., 2017). Chance level is 0.2.

**Table S4.** Results of parameters for metric and cross-validation using all features

| Cross-Validation | Features (All/Best) | Metric | Penalty parameter C (Median) | Penalty parameter C (Std) | Testing Accuracy (Median) | Testing Accuracy (Std) | Training Accuracy (Median) | Training Accuracy (Std) |
| --- | --- | --- | --- | --- | --- | --- | --- | --- |
| WS | AF | BOLD | 0.29 | 0.86 | 0.19 | 0.04 | 0.25 | 0.03 |
| BS | AF | BOLD | 8.64 | 30.31 | 0.24 | 0.02 | 0.26 | 0.00 |
| WS | AF | Cov20 | 1012.72 | 1907.48 | 0.68 | 0.03 | 1 | 0 |
| BS | AF | Cov20 | 0.00 | 0.00 | 0.50 | 0.07 | 0.81 | 0.1 |
| WS | AF | Cov40 | 106.12 | 35.35 | 0.92 | 0.03 | 1 | 0 |
| BS | AF | Cov40 | 592400.6 | 2079076 | 0.60 | 0.09 | 1 | 0 |
| WS | AF | Cov60 | 59.82 | 58.09 | 0.99 | 0.01 | 1 | 0 |
| BS | AF | Cov60 | 591719.8 | 2079269 | 0.69 | 0.1 | 1 | 0 |
| WS | AF | Cov80 | 94.33 | 47.15 | 0.99 | 0.01 | 1 | 0 |
| BS | AF | Cov80 | 1155225 | 2829641 | 0.62 | 0.14 | 1 | 0 |
| WS | AF | Cov100 | 70.74 | 57.76 | 1 | 0 | 1 | 0 |
| BS | AF | Cov100 | 577996 | 2082457 | 0.77 | 0.11 | 1 | 0 |
| WS | AF | Cov120 | 24.17 | 46.88 | 1 | 0 | 1 | 0 |
| BS | AF | Cov120 | 757.46 | 1661.84 | 0.77 | 0.12 | 1 | 0 |
| BS | AF | sCov | 14186.89 | 50856.45 | 0.7 | 0.23 | 1 | 0.07 |
| WS | AF | Covvar20 | 530.66 | 1432.91 | 0.62 | 0.04 | 1 | 0 |
| BS | AF | Covvar20 | 2324511 | 3644483 | 0.44 | 0.08 | 0.99 | 0.01 |
| WS | AF | Covvar40 | 577.24 | 1416.75 | 0.88 | 0.03 | 1 | 0 |
| BS | AF | Covvar40 | 591795.2 | 2079248 | 0.53 | 0.09 | 1 | 0.00 |
| WS | AF | Covvar60 | 117.91 | 0 | 0.97 | 0.01 | 1 | 0 |
| BS | AF | Covvar60 | 2325932 | 3643578 | 0.6 | 0.12 | 1 | 0 |
| WS | AF | Covvar80 | 94.33 | 47.15 | 0.99 | 0.01 | 1 | 0 |
| BS | AF | Covvar80 | 1155621 | 2829479 | 0.62 | 0.13 | 1 | 0 |
| WS | AF | Covvar100 | 59.82 | 58.09 | 1 | 0.00 | 1 | 0 |
| BS | AF | Covvar100 | 578701.7 | 2082262 | 0.64 | 0.14 | 1 | 0 |
| WS | AF | Covvar120 | 71.05 | 57.40 | 1 | 0.00 | 1 | 0 |
| BS | AF | Covvar120 | 578365.5 | 2082355 | 0.63 | 0.14 | 1 | 0 |
| WS | AF | PC20 | 59.55 | 58.36 | 0.66 | 0.03 | 1 | 0 |
| BS | AF | PC20 | 14455.62 | 50797.02 | 0.43 | 0.07 | 0.99 | 0.00 |
| WS | AF | PC40 | 117.91 | 0 | 0.90 | 0.02 | 1 | 0 |
| BS | AF | PC40 | 14119.31 | 50875.18 | 0.54 | 0.11 | 1 | 0.00 |
| WS | AF | PC60 | 83.12 | 53.15 | 0.98 | 0.01 | 1 | 0 |
| BS | AF | PC60 | 42.12 | 56.49 | 0.60 | 0.13 | 1 | 0.00 |
| WS | AF | PC80 | 106.40 | 34.51 | 1 | 0.00 | 1 | 0 |
| BS | AF | PC80 | 16.89 | 41.24 | 0.61 | 0.14 | 1 | 0 |
| WS | AF | PC100 | 82.82 | 53.60 | 1 | 0.00 | 1 | 0 |
| BS | AF | PC100 | 8.67 | 30.31 | 0.64 | 0.15 | 1 | 0 |
| WS | AF | PC120 | 48.03 | 57.06 | 1 | 0.00 | 1 | 0 |
| BS | AF | PC120 | 14144.6 | 50868.18 | 0.66 | 0.18 | 1 | 0.01 |
| BS | AF | sPC | 362.21 | 1238.77 | 0.8 | 0.18 | 1 | 0 |
| WS | AF | GL Prec20 | 0.29 | 0.86 | 0.68 | 0.03 | 0.99 | 0.00 |
| BS | AF | GL Prec20 | 0.00 | 0 | 0.53 | 0.11 | 0.99 | 0.00 |
| WS | AF | GL Prec40 | 518.31 | 1436.94 | 0.88 | 0.02 | 1 | 0.00 |
| BS | AF | GL Prec40 | 0.02 | 0.03 | 0.66 | 0.13 | 1 | 0.00 |
| WS | AF | GL Prec60 | 553.95 | 1425.04 | 0.97 | 0.01 | 1 | 0 |
| BS | AF | GL Prec60 | 93.26 | 47.19 | 0.73 | 0.16 | 1 | 0 |
| WS | AF | GL Prec80 | 554.23 | 1424.93 | 1 | 0.01 | 1 | 0 |
| BS | AF | GL Prec80 | 51.57 | 57.45 | 0.76 | 0.17 | 1 | 0 |
| WS | AF | GL Prec100 | 518.58 | 1436.84 | 1 | 0.00 | 1 | 0 |
| BS | AF | GL Prec100 | 59.99 | 57.93 | 0.82 | 0.17 | 1 | 0 |
| WS | AF | GL Prec120 | 24.45 | 46.74 | 1 | 0 | 1 | 0 |
| BS | AF | GL Prec120 | 14498.76 | 50784.77 | 0.86 | 0.19 | 1 | 0 |
| BS | AF | GL sPrec | 14464.69 | 50794.44 | 1 | 0.20 | 1 | 0 |
| WS | AF | GL Cov20 | 12.13 | 35.27 | 0.70 | 0.04 | 1 | 0.00 |
| BS | AF | GL Cov20 | 690.13 | 1688.56 | 0.55 | 0.12 | 1 | 0.00 |
| WS | AF | GL Cov40 | 94.90 | 46.01 | 0.86 | 0.03 | 1 | 0.00 |
| BS | AF | GL Cov40 | 1413.45 | 2158.98 | 0.63 | 0.14 | 1 | 0.00 |
| WS | AF | GL Cov60 | 94.90 | 46.01 | 0.95 | 0.02 | 1 | 0 |
| BS | AF | GL Cov60 | 117.91 | 0 | 0.70 | 0.14 | 1 | 0 |
| WS | AF | GL Cov80 | 106.40 | 34.51 | 0.99 | 0.01 | 1 | 0 |
| BS | AF | GL Cov80 | 413.14 | 1225.16 | 0.69 | 0.15 | 1 | 0 |
| WS | AF | GL Cov100 | 59.55 | 58.36 | 1 | 0.01 | 1 | 0 |
| BS | AF | GL Cov100 | 577651.8<br>5 | 2082552.3<br>4 | 0.68 | 0.14 | 1 | 0 |
| WS | AF | GL Cov120 | 47.47 | 57.52 | 1 | 0.00 | 1 | 0 |
| BS | AF | GL Cov120 | 43.96 | 55.12 | 0.67 | 0.14 | 1 | 0 |
| BS | AF | GL sCov | 380.07 | 1234.21 | 0.7 | 0.23 | 1 | 0 |
| WS | AF | iPC | 0.87 | 1.32 | 0.42 | 0.03 | 0.84 | 0.09 |
| BS | AF | iPC | 0.00 | 0.00 | 0.33 | 0.04 | 0.75 | 0.08 |
| WS | AF | eigPC | 808640.7 | 2425920 | 0.32 | 0.02 | 0.74 | 0.12 |
| BS | AF | eigPC | 577600.3 | 2082567 | 0.27 | 0.03 | 0.63 | 0.00 |

\* Rounded to two decimals.

**Table S5.**
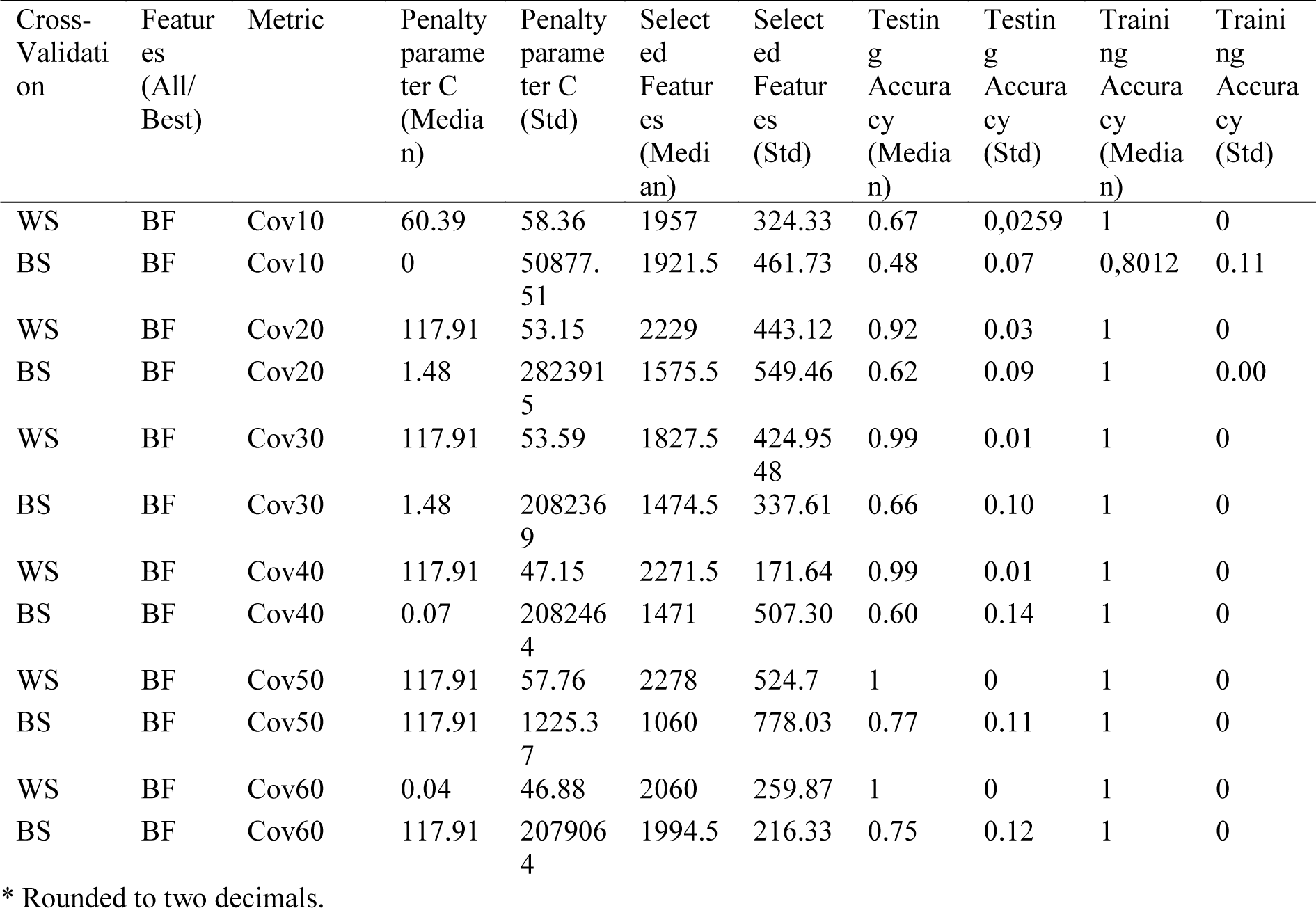
Results of parameters for metric and cross-validation using best features

